# Data-driven re-referencing of intracranial EEG based on independent component analysis (ICA)

**DOI:** 10.1101/150045

**Authors:** Sebastian Michelmann, Matthias S. Treder, Benjamin Griffiths, Casper Kerrén, Frédéric Roux, Maria Wimber, David Rollings, Vijay Sawlani, Ramesh Chelvarajah, Stephanie Gollwitzer, Gernot Kreiselmeyer, Hajo Hamer, Howard Bowman, Bernhard Staresina, Simon Hanslmayr

## Abstract

Intracranial recordings from patients implanted with depth electrodes are a valuable source of information in cognitive neuroscience. They allow for the unique opportunity to record brain activity with a high spatial and temporal resolution. To extract the local signal of interest in stereotactic EEG (S-EEG) data, a common pre-processing choice is to re-reference the data with a bipolar montage.

With bipolar reference, each channel is subtracted from its neighbour in order to reduce commonalities between channels and isolate activity that is spatially confined. We here challenge the assumption that bipolar reference can effectively perform this task. We argue that in order to extract local activity, the distribution of the signal source of interest, as well as the distribution of interfering distant signals and noise sources need to be considered. Those can have a variable spatial extent and are modulated by electrode spacing, location and anatomical characteristics. Those factors are not accounted for by a fixed referencing scheme and bipolar reference can therefore not only decrease the signal to noise ratio (SNR) of the data, but also lead to mislocalization of activity and consequently to misinterpretation of results.

We promote the perspective of regarding referencing as a spatial filtering operation with fixed coefficients. As an alternative, we propose to use Independent Component Analysis (ICA), to derive filter coefficients that reflect the statistical dependencies of the data at hand. We argue that ICA performs the same task that bipolar referencing pursues, namely undoing the linear superposition of activity and can therefore be used to identify activity that is local. We first describe and demonstrate this procedure on human S-EEG recordings. In a simulation with real data, we then quantitatively show that ICA outperforms the bipolar referencing operation in sensitivity and importantly in specificity when it comes to revealing local time series from the superposition of neighbouring channels.

## Introduction

An increasingly popular tool in modern cognitive neuroscience is to investigate local field potentials recorded from electrodes implanted into the brain in epileptic patients. Data recorded from these patients offer a unique and powerful opportunity to understand how local neural populations implement cognitive operations. As is the case in all EEG recordings electrical potentials are recorded as differences between two sites, in the simplest case an “active” site and a “passive” reference. Therefore a delicate decision has to be made as to what is the best reference for a give recording site (e.g. Nunez and Srinivasan, 2006). While online recording of stereotactic EEG (S-EEG) can for example be performed with a mastoid or a subdermal reference, the offline analysis is often preceded by a re-referencing operation. It is a popular choice to re-reference with a bipolar montage (e.g. Staudigl et al., 2015; Tallon-Baudry et al., 2001), in which each channel is subtracted from its neighbour. Bipolar referencing is seen as advantageous since it removes contamination from activity at the online reference electrode, and highlights activity that is local (Lachaux et al., 2003).

In a recent paper Mercier et al. (Mercier et al., 2017) investigate the diffusion of cortical field potential in S-EEG and empirically compare the effect of commonly used reference choices on the recorded signal. They find that using both neighboring electrodes as a reference in a local referencing scheme, reduces correlations between channels. Based on this reference, the authors analyze the activity recorded in cortical white matter and conclude that it contains a mixture of signal sources spreading from nearby gray matter and signal from other sources that sometimes correlates with distant gray matter activity. Their findings highlight the importance of considering anatomical structure when choosing the reference. They further render the use of a white matter reference somewhat suboptimal and put a spotlight on the fact that recorded activity at a given electrode reflects a mixture of activity from different sources.

The authors further highlight that *“re-referencing electrophysiological data is a critical preprocessing choice that could drastically impact signal content and consequently the results of any given analysis”* (Mercier et al., 2017 p. 219, l.4). We fully agree with this statement and think that the problem of referencing needs more discussion. Especially when differential functions of local structures are investigated with S-EEG, a wrong choice of reference can lead to drastic misinterpretations of results; that is, mislocalization of effects. This is highly relevant in cognitive neuroscience, since results from intracranial recordings act somewhat as a ground truth for the localization of electrophysiological activity.

In the current paper, we therefore want to add to this discussion. Specifically, we believe that it is helpful to point out disadvantages of bipolar and local referencing schemes, especially, with regard to activity recorded in subcortical structures (notably the Hippocampus), which were not considered by Mercier et al., due to “*distinct characteristics of the corresponding signal*” (Mercier et al., 2017 p. 221, l. 21). By spelling out the mathematical details of the referencing operation, we more explicitly discuss its effects on signal-to-noise-ratio (SNR) and identify conditions in which bipolar referencing can lead to mislocalization of signal activity. As a potential alternative, we propose to use Independent Component Analysis (ICA) in order to derive a data-driven referencing scheme that is based on the statistical dependencies between channels. In this context, we champion a different perspective on referencing schemes by describing them as spatial filtering operations. For simplicity, in our comparisons, we will mainly focus on the bipolar referencing scheme that is often the preferred preprocessing choice; however most of our analysis and simulations also generalize to the local referencing scheme proposed by Mercier et al. (Mercier et al., 2017).

## Material and Methods

### Participants

For the analyses, data from 3 female patients were used who were 24, 33 and 42 years old. Two patients were recorded at the Queen Elizabeth Hospital Birmingham (QEHB), 1 patient was recorded in University Hospital Erlangen (UKE).

Patients suffered from drug-resistant epilepsy and were implanted with intracranial depth electrodes. They underwent pre-surgical monitoring purely for diagnostic purposes. All patients volunteered to participate in a memory study and formed part of a larger sample of volunteers. Written informed consent was obtained in accordance with the Declaration of Helsinki.

### Task

During the memory task, patients were watching 3 different 4-second-long movie sequences that each consisted of 2 distinct scenes. In one of the scenes, a word appeared in the center of the screen and patients were instructed to vividly associate the word with the exact scene within the movie. After performing a short distractor task in which they categorized numbers as odd or even, patients were shown the words in an arbitrary order. For every word they first decided whether they had previously learned it in scene 1 or scene 2 of a movie, and then identified the movie it was associated with. To improve performance and increase the number of successfully remembered trials, each association was learned 3 times and later recalled 3 times.

### Data recording and preprocessing

The data was continuously recorded at a sampling rate of 1024 Hz with an online linked-mastoid reference. Data was imported into MATLAB 2014a (MathWorks) using FieldTrip (Oostenveld et al., 2011) for data from QEHB and Brainstorm (Tadel et al., 2011) for data from UKE. Subsequently, data were downsampled to 1000 Hz and epochs of 7 seconds were created, beginning 2 seconds before video onset at encoding and word onset at retrieval. Epochs ended 5 second after video/word onset.

All channels that displayed frequent epileptic spiking or electrical noise were excluded from further analysis. Remaining trials that still contained artifacts were manually removed at a later stage.

### ICA computation

ICA was computed using the EEGLab implementation ‘runica’ (Delorme and Makeig, 2004). Before computation of the unmixing matrix, all data was filtered with a high-pass filter of 1.5 Hz and trials containing artifacts were heuristically removed based on statistical characteristics (i.e. variance, kurtosis and maximum value). The unmixing-matrix was later applied to the unfiltered data including all trials; trials containing artifacts were then removed based on visual inspection. For the illustration in Fig. 1, only those trials that remained after visual inspection were included and ICA computation was limited to the three channels displayed in the figure.

**Fig. 1.**
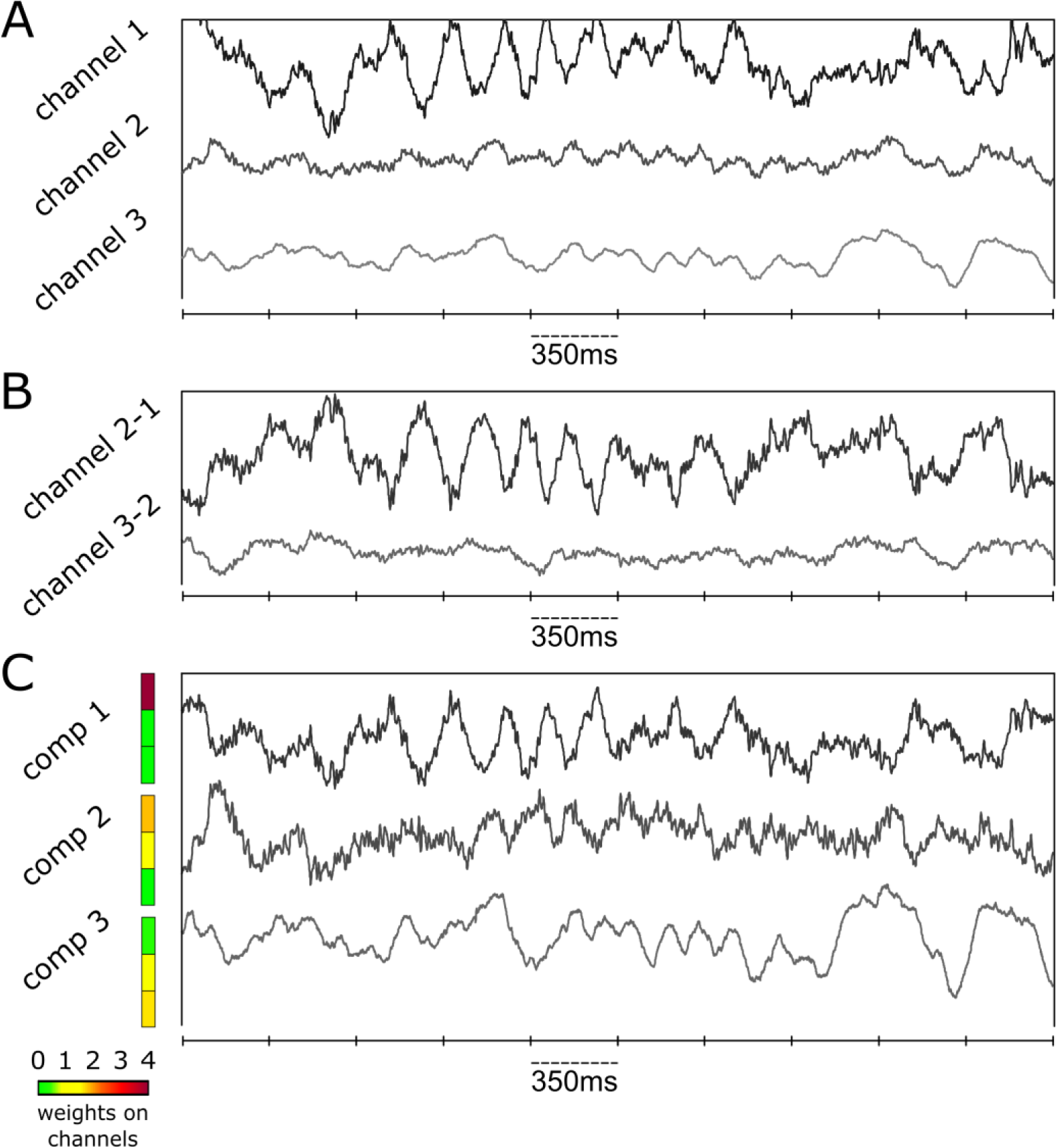
Example of how bipolar re-referencing can change the data. A: Activity during a single trial on 3 selected channels in the medial temporal lobe of a patient implanted with intracranial electrodes. Channels 1 and 2 fall in the Hippocampus, channel 3 is located in the nearby white matter. B: The same channels were re-referenced with a bipolar scheme. The first channel appears inverted and the shared activity between channel 2 and 3 is almost cancelled out. C: The same trial using an ICA decomposition of the data. Inspecting the weights in the columns of the 3x3 mixing matrix (i.e. the topography), we can see how the hidden components mix into the channels observed in A. Instead of subtracting the channels from each other and therefore mixing them even more, ICA splits the 3 channels into activity that is unique to channel 1 (component 1), activity that is shared between channel 1 and 2 (component 2) and activity that is shared between channel 2 and 3 (component 3). Interestingly component 2 is mostly present in channel 1, however it appears strongest (and inverted) on the combined bipolar channel 3-2. This is because the difference between its contribution to channel 1 and to channel 2 is smaller than the difference between its contribution to 2 and to 3, which can be observed in C.

### Electrode localization

For the example-channels displayed in Fig. 1 and Fig. 2, the pre-surgical MRI was first segmented using Freesurfer (Fischl, 2012; Reuter et al., 2012) and anatomical labels were derived from the Desikan-Killiany Atlas (Desikan et al., 2006). Electrode locations were manually determined based on the center of signal drop-out in the post-surgical MRI-scan.

**Fig. 2.**
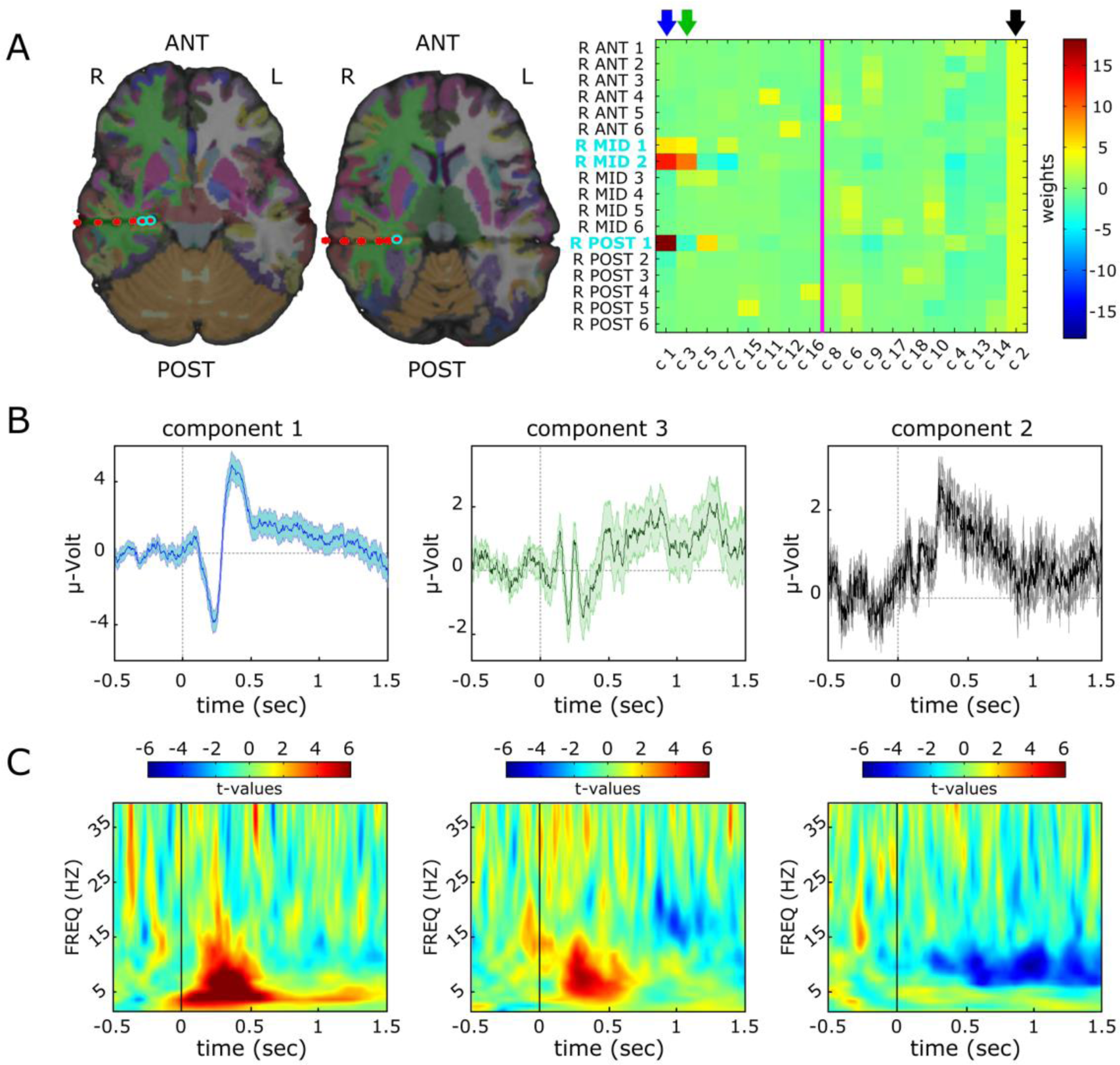
Example of ICA-components and their spatial distribution in one patient with intracranial electrodes. The patient was performing a memory task in which they repeatedly associated short video clips with words and recalled them later. A: two out of three used electrode shafts are depicted on the left. The brain is superimposed by labels corresponding to the Desikan-Killiany Atlas, which were derived via Freesurfer parcellation. R-Mid 1-6 and R-Post 1-6 are located in the right medial and right posterior temporal lobe respectively (counting from inside to outside). ICA was computed and the inverse of the unmixing-matrix (i.e. the topography) was sorted by the spatial “broadness” of the components. Columns of this mixing-matrix (right) correspond to the distribution of the components. The pink vertical line separates local components on the left from components that have a broad distribution across channels (corresponding to a *chi^2^*- value of *p* > 0.2), which one could consider discarding from further analysis. Importantly, the second component (explaining the second most projected variance in the dataset) has an almost uniform distribution across channels, i.e. it is mixed into every channel to an equal extend. This component probably picks up activity from the mastoid-reference. The electrodes labelled in turquois (R-Post 1, R-Mid 1-2) fall inside the right Hippocampus (R-Mid 3 falls partly in the nearby white matter). Component 1 and 3 are independent sources that each peak inside the Hippocampus, with activity related to component 1 originating from the posterior HC, even though it is picked up strongly on R-Mid 2 and R-Mid 1 as well. B-C: Evoked responses and standard error (B) of component 1-3, during the first 1.5 seconds of the movie clips and event locked changes in power spectral density (PSD) during this time (C). Evoked responses were baseline corrected to -500 to -100 ms before the onset of the movie. To show changes in PSD, the power-spectra were rank transformed across 192 trials and a dependent-sample t-test was computed between the PSD rank at each time point and the average PSD rank between -500 and -100 ms. Interestingly, all components show event related changes and the profiles of the broad component and the HC-sources have very distinct properties.

The post-surgical MRI-scan was then co-registered with the pre-operational MRI using robust coregistration (Reuter et al., 2010) as implemented in Freesurfer. Lastly, the labeled segmentation was overlaid with the post-operational MRI and the highlighted electrode positions to determine the approximate structure in which the electrode was located. This was done to derive labeling in a partly automated and standardized way.

The locations of the three electrodes for simulations were manually labeled based on the anatomical characteristics of the surrounding tissue in the post-operational MRI.

### Power Spectra

To derive the power spectra in the encoding block, Fourier transformed data was multiplied with a Hanning taper of 4 cycles for a given frequency. Power spectra of every full frequency between 2 and 39 Hz were computed. This was done every 10ms in a sliding window from 0.5 seconds prior to movie onset to 1.5 seconds during the movie (e.g. Jokisch and Jensen, 2007).

All processing of the data was done using the FieldTrip toolbox for EEG/MEG-analysis (Oostenveld et al., 2011).

### Simulations

For all simulations, three electrodes from three different patients were selected, which were later used to define independent sources based on real data. Electrodes were located in the right Parahippocampal Cortex, in the right Perirhinal Cortex and in the left Middle Temporal Gyrus. For the simulations, ICA was run on unfiltered visually inspected data. After Fisher Z-transformation of correlation coefficients, a dependent samples t-test was computed to derive analytical statistics across 100 repetitions of the simulation.

## Bipolar and local (re-)referencing

Bipolar referencing is a popular preprocessing choice for several reasons (Lachaux et al., 2003; Shirhatti et al., 2016; Trongnetrpunya et al., 2015):

Firstly, bipolar re-referencing removes the reference activity. Activity from the online reference channel is expressed on all channels, since it is subtracted into all electrodes during recording. By later re-referencing one channel against another, the reference activity is removed.

Secondly, bipolar referencing removes noise sources with a broad spatial distribution; if all electrodes are picking up an external noise source (e.g. 50Hz line noise), this noise will likewise disappear in the subtraction of a channel from its neighbor. In practice however, line noise and other noise sources often do not affect all electrodes to the same extent. Therefore, if the data is not sufficiently inspected before re-referencing, this can for example lead to the subtraction of noise into an otherwise clean channel; in other words, the distribution of external noise sources in the data needs to be considered.

Finally, bipolar referencing emphasizes spatially local activity. By subtracting one channel from its neighbor, only signal that is unique to this channel will supposedly remain unaffected. Common (broad) activity is attenuated or even fully removed. This postulation is somewhat problematic, since it relies on a number of assumptions (see below). For the extraction of local signal, the distribution of the signal source of interest, as well as the distribution of interfering distant signals and noise needs to be considered, which can have a variable spatial extent (Kajikawa and Schroeder, 2011).

### The referencing operation

In general, a referenced channel, whether it was computed via bipolar, local, or average referencing will be a simple linear combination of electrodes and therefore a mixture of activity recorded from several electrodes. Whether this linear combination reduces the dependencies between the channels or even introduces new dependencies and mixes separate sources even more, depends on the spatial location and extent of the underlying signal of interest, as well as factors like noise distribution, online reference and electrode location. While the importance of considering anatomical structure in the reference choice has been extensively analyzed by Mercier et al.

(Mercier et al., 2017), we want to consider the implicit assumptions about the distribution of underlying source-signals in bipolar referencing and their impact on the resulting channel-activity.

In the bipolar referencing scheme, the new re-referenced channel is a linear mixture of two electrodes: the time series on the neighboring channel is subtracted from the time series on the electrode of interest.

Using one neighbor as a reference (e.g. Staudigl et al., 2015; Tallon-Baudry et al., 2001), we define the bipolar reference in accordance with the nomenclature proposed by Mercier el al. (Mercier et al., 2017) (eq. 1) as
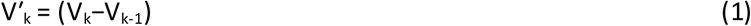

where V_k_ refers to the electrode in the kth position and V’_k_ is the re-referenced time series.

Considering the time series on electrode V_k_, we can write it as a linear combination of activity that we would take for signal S_k_ and activity that is noise N_k_:
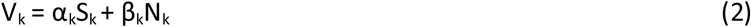

Here, we conceive of noise on a given electrode V, as both true sensor noise and general brain activity that does not emanate from the immediate vicinity of the electrode of interest; that is, brain activity either spread through passive conduction or transported via white matter tracts. If we now compute
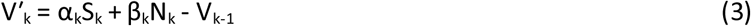

we implicitly expect that activity that is shared between the neighboring channels will constitute mostly noise and that the channel V_k-1_ contributes little or no unique activity to the re-referenced signal. We now want to define S^*^_k-1_ as signal that is unique to contact (k-1) and N^*^_k-1_ as noise that is unique to contact (k-1). Together they account for the remaining activity that is not shared with electrode V_k_.

Since we defined V_k_ as the electrode of interest, the activity at electrode V_k-1_ would be written as
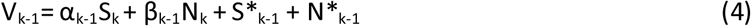

that is, a combination of activity S that would be considered signal at the electrode V_k_, activity N that would be considered Noise at the electrode V_k_, activity S^*^_k-1_ which is unique to the electrode V_k-1_ and would be considered part of the signal at V_k-1_ and noise N^*^_k-1_ which is unique to the electrode V_k-1_. Combining these terms with the previous formula (3), we obtain the activity at the new re-referenced channel V′_k_ as
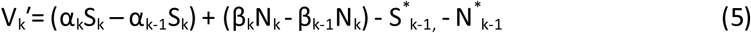

Consequently, the best conditions for bipolar referencing are, if the common noise term Nk is of equal magnitude on both channels (β_k_ = β_k-1_), so it can cancel out, and if the local signal of interest S_k_ is considerably stronger on the contact V_k_ than on the contact V_k-1_ (α_k_ ≫ α_k-1_) and therefore not dampened much. Additionally, little or no unique signal S* _k-1_ and unique noise N*_k-1_ should be present on the neighboring channel V_k-1_ which could otherwise overshadow the signal S_k_.

In practice, we accept that the re-referenced channel V′_k_ will consist of a linear mixture of dampened signal and residual noise from V_k_ and inverted unique signal and unique noise from V_k-1_.

After bipolar referencing, the spatial resolution of the data will be reduced: we can maximally locate the signal of interest between the two electrodes V_k_ and V_k-1_. However, a signal of interest emanating in the close vicinity of V_k_ or V_k-1_, will not automatically have its maximal strength between these electrodes after re-referencing. Since subtracting one channel from its neighbor is essentially taking the spatial derivative, the strongest signal after re-referencing will appear at the point where the signal drops/increases most between two channels. This means that for a correct localization, the difference in signal strength α between electrode k and (k-1) must be bigger than between (k-1) and (k-2) and bigger than between (k+1) and k (i.e. |α_k+1_ – α_k_ |<| α_k_ – α_k-1_ |>|α_k-1_ – α_k-2_|).

This requirement can be especially problematic in cases where some electrodes on a shaft are in a discrete structure. The two most mesial electrodes on a shaft can for example fall in the Hippocampus and pick up a strong signal there. A third electrode in the Parahippocampal Cortex can pick up very little of this activity. In the re-referenced channels, it will then appear like the peak of the underlying hippocampal signal is between Hippocampus and Parahippocampal Cortex (or incorrectly interpreted on the re-referenced channel V’_3_ (= V_3_-V_2_) which falls in the Parahippocampal Cortex), even though only the two electrodes in the Hippocampus strongly picked up the characteristic signal in the first place.

In another hypothetical scenario, consider a signal that has a very broad distribution and is nearly equally strong on the first (n-1) electrodes of the shaft. It will only appear in inverted form at the end of the electrode shaft after bipolar re-referencing and could easily be mistaken for activity from a local source. These are just two examples of how the standard approach of bipolar referencing can lead to drastic mislocalization of a source signal.

### Impact on SNR

An important quantity for assessing signal quality after re-referencing is signal-to-noise-ratio (SNR). In the bipolar referencing scheme, SNR can either increase or decrease, depending on the distribution of signal of interest (S), the distribution of signal that is not of interest (S*) and the distribution of noise sources (N and N*).

If we quantify the signal to noise ratio before re-referencing as
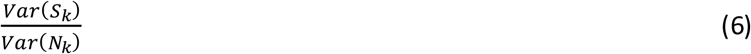

we can write the signal to noise ratio of the re-referenced channel as
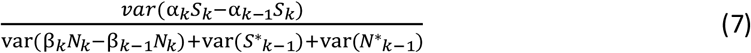

for which we used that the unique signal S*_k-1_ and the unique noise N*_k-1_ are by definition independent of each other and independent of the signal and noise time series S_k_ and N_k_.

Generally, only noise that is shared between a selected channel and the neighboring channel will have a positive impact on SNR with re-referencing. Shared signal will lead to a dampening of the signal, i.e. decrease the numerator in the Signal/Noise fraction and therefore decrease the signal to noise ratio on the re-referenced electrode, whereas unique noise and signal that is not of interest (e.g. stemming from white matter) will add to its denominator. Under favorable conditions, the bipolar referencing scheme can therefore improve SNR, however it can also decrease SNR and signals can appear stronger on neighboring or even distant channels than on the source-channel (see above).

Notably, reduction in SNR will also decrease correlations between channels i.e. a reduction in correlation as observed by Mercier et al. (Mercier et al., 2017), can occur partly because signal is lost in the re-referencing process. In other words: noise can be uncorrelated as well, so a lack of correlation alone is not a reliable quality measure for successful extraction of a local signal of interest.

### Referencing is spatial filtering

A useful perspective in assessing referencing schemes is to consider them as spatial filters. In analogy to temporal filters, spatial filters can sometimes be characterized by the spatial frequencies that are selected or suppressed.

The bipolar referencing operation is an approximation of the first spatial derivative. Since the derivative and the Fourier transform are both linear operations, we can estimate the effect of bipolar referencing on the spatial frequency spectrum by taking the derivative of the Fourier-coefficients, in which each coefficient’s complex conjugate is multiplied by the frequency itself. This means that high frequencies are amplified and low frequencies are dampened.

If we assume that low temporal frequencies will show a broader spread to neighboring electrodes than high temporal frequencies (i.e. also have a lower spatial frequency), the shared signal between neighboring electrodes will be stronger in the low frequencies and therefore low temporal frequencies will be disproportionally dampened by the referencing operation, in other words the spectral properties of the time series can change.

Importantly, these confounds will be modulated by the distance between neighboring electrodes and their location (i.e. Manufacturers offer a variety of models with different spacing between electrodes), impeding the comparability between studies and even patients.

Furthermore, a general problem with local/bipolar referencing is the inevitable loss of information. Activity on the *n* electrodes on an electrode shaft cannot be sufficiently explained by the linear combination of (n-1) electrodes that is obtained via bipolar referencing ((n-2) electrodes in local referencing) unless there are already linear dependencies between the channels beforehand (i.e. the data is rank deficient). This problem is aggravated in datasets in which only few electrodes are present on each shaft.

### Towards a data driven reference

The purpose of this manuscript is to demonstrate that ICA (Comon, 1994; Hyvärinen and Oja, 2000) outperforms bipolar referencing (and other referencing schemes) in extracting the signal of interest. This applies specifically to cases in which the goal of referencing is to extract sources that are local and have high spatial specificity (i.e. do not pick up activity that originates in distant structures).

The goal of ICA is very similar to that of bipolar referencing, namely to apply spatial filters to the recorded activity in order to undo the linear superposition of sources and find the underlying signal time series.

By separating the activity into the same number of sources as there are channels, no information is lost in this process. The filters that are computed via ICA not only optimize the statistical independence between underlying components (which is a stronger requirement than merely reducing correlations (Rodgers et al., 1984)), they also allow for hidden sources with high and low spatial frequency and this distribution can be inspected. The resulting filter coefficients will be static across time and ICA therefore acts in the same way as referencing operations: each channel is replaced with a linear combination of channels.

In contrast to the classical referencing schemes, ICA is a data-driven approach. That is, its coefficients adapt to the data at hand, whereas bipolar referencing uses the fixed coefficients (1,-1) on neighboring channels. Thus bipolar referencing is but one implementation of a much larger space of coefficients afforded by ICA.

## Proposition: ICA can detect local sources

Independent component analysis (ICA) is a method for blind source separation. It aims at explaining a random vector by a linear combination of underlying components (sources) that are statistically independent (Comon, 1994).

ICA is extensively used in scalp EEG/MEG in order to separate brain signals from artifacts (Delorme and Makeig, 2004).

In the underlying framework, the observed data can be sufficiently explained by a linear mixture of underlying sources:
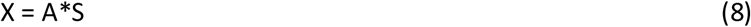

where X is the data (channels x time points), S are the underlying sources (components x time points) and A is a transformation matrix, which is usually called the mixing matrix.

In practice, the mixing matrix A and the underlying sources are both unknown. ICA aims to undo the linear mixing – algorithms find the inverse of the mixing matrix A^-1^ (i.e. the unmixing matrix), such that the resulting components are maximally statistically independent:
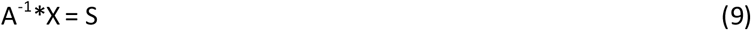

See (Hyvärinen and Oja, 2000) for a review. Since A^-1^ is a square matrix of full rank, it is invertible and we can switch between channel representation and component representation of the data without losing information. Importantly, the rows of the unmixing matrix act as a set of spatial filters, i.e. they are linear transformations of the electrodes that highlight the activity of an independent source. When we inspect the columns of the mixing matrix A, we can see how much an underlying source contributes to each channel in the observed data representation.

Usually, a column of the mixing matrix can be used to visualize the topography of a component, which is helpful in detecting artifacts in scalp M/EEG (Delorme and Makeig, 2004). For intracranial data, we can still use this information in order to identify local sources and separate them from hidden sources that affect all channels to an almost identical extent.

Intuitively, ICA feels like a drastic transformation of the data and it appears less justifiable then simple re-referencing. In particular, the ICA framework relies on assumptions like a non-Gaussian distribution of underlying sources and a linear mixing model. In practice, however, the output of the ICA is a set of linear filters that optimally complies with these assumptions and unmixes the linear superposition on the channels accordingly. The strong benefit from this is that no a priori assumptions about the filter-coefficients are made.

If the goal of the analysis is to draw conclusions about local sources, we therefore propose to do an ICA on the data and then compute a measure of uniformity for every column in the mixing matrix. This has the advantage that no prior assumptions about the distribution of signal and noise on the electrodes are made; rather the actual signal spread is estimated from the statistical properties of the data. Later, the distribution of underlying sources can be inspected and broad sources can be discarded from further analysis.

In particular, we propose the following steps:

1) Compute ICA on a cleaned version of the data, excluding channels and trials that are contaminated by epileptic spikes.
2) Calculate a measure of uniformity (‘broadness’) for the absolute (since a component can have positive and negative polarity on different channels) of every column of the mixing matrix. As a measure of uniformity, we propose a simple chi^2^-value; degrees of freedom correspond to the number of channels. Alternatively, one could consider the Kullback–Leibler divergence (Kullback and Leibler, 1951) from a uniform distribution or a related measure.
3) Discard all sources that are broad in topography (e.g. the column has a chi^2^ value higher than a threshold of p > 0.2) from further analysis.
4) One of a, b, c:
  a) Project the remaining components back in order to keep working on channel-data. The channels in the cleaned dataset will have linear dependencies; effects can be interpreted as being present in the structure they are measured in and not originating from broad sources or the reference channel.
  b) Continue working with the local independent components. Since the largest weight in the column of the mixing matrix determines which channel picks up most of the activity from that component, component-labels can be changed to the label of their peak-weight. The reasoning behind this is that signals should be strongest where they originate. Additionally, all other weights can be kept. They can be useful when interpreting the spatial extent of an effect and when averaging across several subjects.
  c) Selectively build channels from a linear combination of only those independent components that have their largest weight (peak in the column of the mixing matrix) on the corresponding electrode. This way, in the linear combination of a channel, only contributions of identified sources at this electrode are considered. This probably produces the most realistic representation of the signal that would be measured by a reference-free, uncontaminated electrode in the respective structure, without interference from distant sources.

An example of steps 1-3 is given in Fig. 2. Data from one patient is presented that had electrodes implanted in the left and right hemisphere. ICA was computed on the channels from the right hemisphere, whereas channels from the left were discarded due to epileptic spiking. Three shafts were located in the right anterior, medial and posterior temporal lobe; some channels on the medial and posterior shaft were located in the Hippocampus (Fig. 2A, left). A chi^2^ test was then computed on the columns of the absolute of the mixing matrix. Columns of the mixing matrix were then sorted by the chi^2^ value in descending order (Fig. 2A, right). The resulting component 1 and component 3 showed a local distribution and peaked in the Hippocampus. It can be seen that component 1 is expressed most strongly on the posterior hippocampal channel R POST 1, but it also affects the channel in the medial Hippocampus to a larger extent than any other source. Another local source could be revealed on the medial hippocampal channel R MID 2. Importantly, the component which explained the second most mean projected variance in the data (i.e. component 2) had an almost uniform distribution. It can therefore not be considered local activity and was probably due to activity on the reference mastoid electrodes. All components 1-3 expressed an evoked response upon stimulus onset (Fig. 2B). They also showed distinct responses in the power spectrum upon stimulus presentation, namely a strong theta power increase was observed in the local hippocampal sources, whereas the broad source showed mostly alpha power decreases, consistent with this source picking up scalp EEG from the mastoids (Fig. 2C). This illustrates the importance of separating local and broad activity when analyzing intracranial EEG and the power of ICA in effectively performing this task.

### Simulation of the influence of spatial frequency and noise on signal recovery in ICA and bipolar reference

To quantitatively compare the abilities of ICA and local referencing in uncovering the activity of local sources, we performed simulations using real iEEG data.

For simplicity, we compared ICA to a bipolar referencing scheme using only three electrodes. However, the observed principles generalize to other setups with a larger number of electrodes.

Firstly, we defined three latent sources. To this end, 3 different channels were selected using datasets from 3 different patients. This allowed us to use real data but avoided signal spread or other unwanted statistical dependencies. Note that using real channel data in isolation will present a challenging scenario for ICA since this recorded activity from each patient is itself a superposition of other sources and will therefore likely violate the assumption of a non-gaussian distribution, which means that in a realistic scenario, ICA should perform a lot better. A total of 480,000 sampling points was now used on each channel, which (at a sampling rate of 1000 Hz) corresponds to an 8-minute recording. These three channels served as source 1, source 2 and the source which acted as underlying reference. All channels were demeaned and scaled to unit variance; the reference source was scaled to 10 % of the strength of the other sources to reflect the fact that in practice reference channels should pick up little activity.

Next we defined a linear mixture model: Sources 1 and 2 were placed on the outer electrodes 1 and 3 (i.e. the source-channels affected these electrodes with strength of 1) and spread into the neighboring electrodes with a varying strength of 1/a and 1/a^2^ respectively (Fig. 3A). The mixing parameter *a* was decreased in 40 logarithmic steps from 10 to 1.02 in order to increase the “broadness” of the underlying sources, such that they affected neighboring channels to a greater extent.

**Fig. 3.**
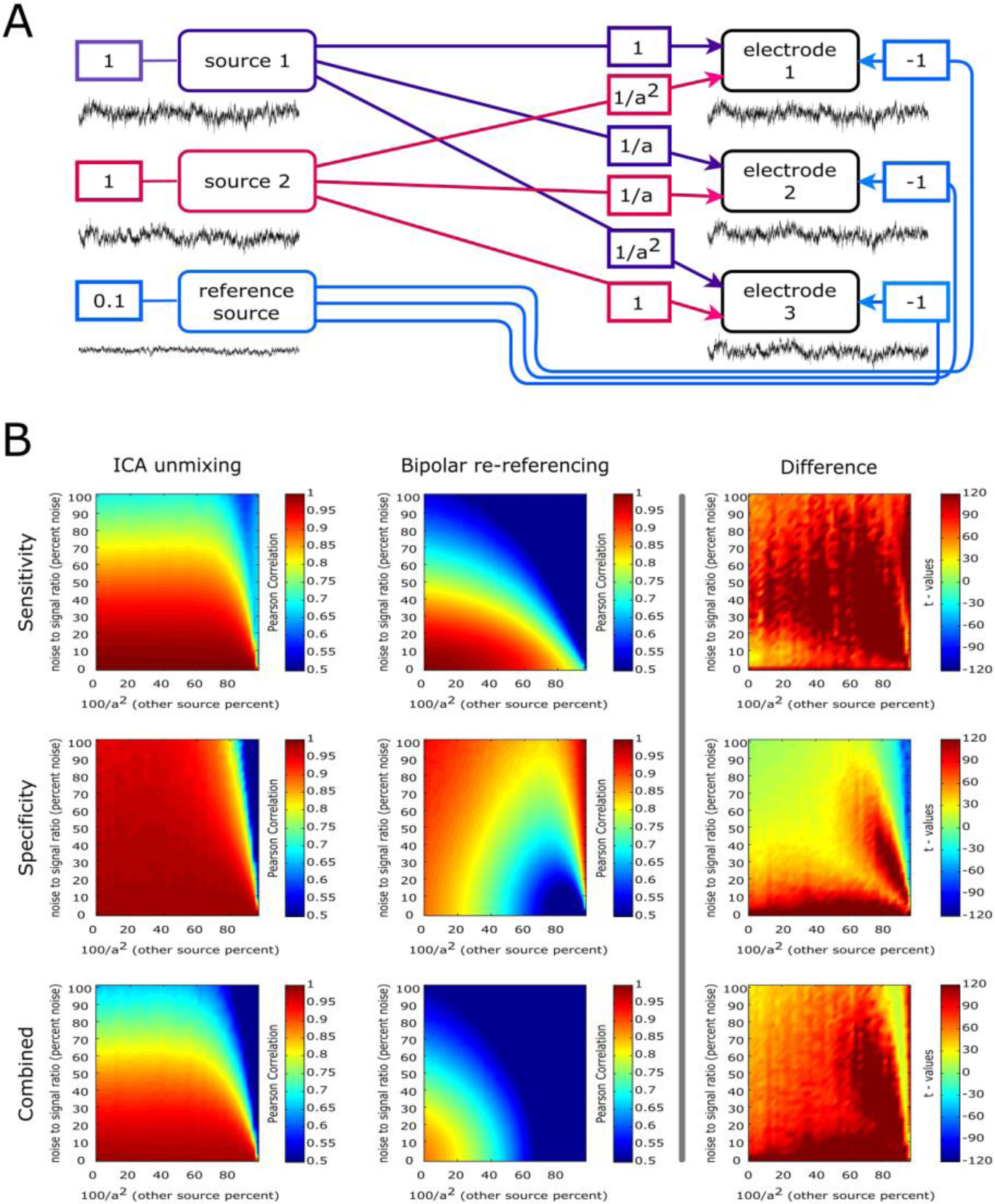
Simulation of the influence of spatial frequency and noise on signal recovery, comparing ICA and bipolar reference. A) A linear mixture model was defined in which two hidden sources and a reference mixed into three neighbouring electrodes. Each source channel was located at one end of the simulated electrode shaft and affected the neighboring electrodes with a decreasing strength of 1/a and 1/a^2^. The factor was decreased in logarithmic steps in order to modulate the “broadness” of the signal spread. B) The recovery performance was evaluated for ICA and bipolar reference for different levels of broadness. Additionally, different levels of noise were simulated by adding independent pink noise of increasing amplitude to the electrodes. In order to determine sensitivity (upper row), the ICA component that peaked on electrode 1 (electrode 3) was correlated with source 1 (source 3). Likewise for bipolar reference, the timecourse of electrode 1-2 (electrode 3-2) was correlated with source 1 (source 2). Specificity (middle row), refers to 1-correlation with the opposite source. The four plots on the top-left show average absolute correlation averaged across both electrodes and 100 repetitions. The two plots below (bottom row) show a combined measure of sensitivity*specificity. The three plots on the right (right column) show the t-statistic of difference acoss 100 runs. ICA showed better performance in recovering the original source (sensitivity) and importantly was more spatially specific than bipolar reference. Bipolar reference performed well when the spatial frequency of underlying sources was high and noise levels were low. The increase of bipolar reference in specificity under high “broadness” of sources and high levels of noise is due to a loss of signal altogether (i.e. noise correlations).

Additionally, the two sources were referenced against the reference signal by subtraction. These linear mixing and referencing operations can be succinctly summarized in the following mixing matrix:
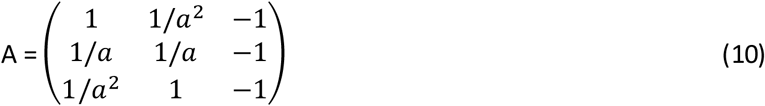

The mixing matrix was then used to define observed data by left-multiplying it with the matrix of source signals as X = A*S, where S are the underlying sources (3 components x time points) with source 1 in the first row, source 2 in the second row and the reference source in the third row. Each row in the mixing matrix specifies how these sources are combined to form the electrode activities. For instance, inspecting row 1 of A, we see that source 1 is added with factor 1, source 2 is added in attenuated form with a factor of 1/a^2^, and the reference is subtracted by adding it with factor -1.

The ICA was now run 100 times on the observed data X for every level of the parameter *a*. Likewise, the bipolar reference was computed by subtracting the second electrode in X (i.e. the second row) from electrode 1 and from electrode 3. We then added increasing levels of pink noise to each individual observed channel in X. In this, we adjusted the variance of the noise signal in 40 linear steps from 0 to 100% of the variance of the signal-sources. Again, we repeated the ICA and the bipolar referencing 100 times for every level of *a* and every level of noise.

We defined the sensitivity of ICA to recover source-channel 1 and 2 respectively, by selecting the recovered components that had their peak weight on one of the outer electrodes; we then took the absolute correlation of each of these components with the respective underlying source signal that had been placed on that channel.

Similarly, we defined sensitivity of bipolar reference by taking the absolute correlation of the re-referenced outer electrodes with their underlying sources.

To define specificity of the recovered source, we computed the correlation with the underlying source that was placed on the opposite end of the three electrodes and used 1 – correlation as a metric. In a combined measure, we then multiplied sensitivity and specificity to account for the fact that specificity can be due to a loss of signal altogether and therefore is only informative in the presence of sensitivity. Average sensitivity, average specificity and the combined measure were compared between ICA and bipolar referencing with a dependent samples t-test of Fisher Z-transformed correlation coefficients.

Results showed that ICA performed constantly better in separating the two sources (Fig. 3B). This was most pronounced when noise levels were high and spatial mixing was high, but interestingly ICA could also recover signals better when noise levels were low and mixing was little. Importantly, a crucial result is that ICA shows constantly better specificity than bipolar referencing. This is a critical point, since the proclaimed goal of bipolar referencing is to extract activity that is local, in other words to be highly spatially specific.

There are some limitations to this simulation. Firstly, we only used an 8-minute recording, whereas in practice an experimental session with a patient may last longer. However, computation on a full recording should result in an even better estimation of the ICA filters, whereas the bipolar reference operation remains the same. Secondly, we only simulated a linear mixture of three channels, whereas in practice a dataset can be formed of up to several hundred electrodes.

## Conclusion

Fixed (re-) referencing schemes such as bipolar reference are standard solutions for preprocessing. They are powerful tools, but they address every dataset in exactly the same way. Given the large variability of S-EEG electrode spacing and location, an adaptive solution seems necessary. Careful preprocessing could entail the hand-picking of reference electrodes based on anatomical structure, however as shown by Mercier et al. (Mercier et al., 2017) even channels located in white matter can contain signal from distant gray matter and re-referencing can therefore lead to the superposition of distant sources in channels.

While local and bipolar referencing schemes are often used to extract signal that is local, we here show that bipolar referencing does not automatically convey spatial specificity, especially when applied without considering anatomical information and the data at hand.

Instead, we propose a data driven approach in which the statistical dependencies between channels are exploited. We want to promote the perspective that referencing is a spatial filtering operation and suggest using ICA, which derives spatial filters that optimally isolate hidden sources.

Hence, we see the spatial filters computed via ICA as a referencing scheme that was derived from the data. In practice, the spatial filters can then actually resemble a bipolar or local referencing scheme, meaning that ICA finds coefficients close to (1, -1) on neighboring electrodes, however the particular advantage of ICA is its adaptiveness; that is, filter coefficients are in no way restricted to prior assumptions. The “broadness” of the mean projected variance across channels should then be inspected for each IC time series in order to exclude (or at least separate) broad activity from the analysis. We propose to use a chi^2^ test in order to quantify the broadness of components, however setting a fixed criterion for broadness can be potentially problematic since the electrode coverage and density will affect those metrics. A potential solution to this could be to incorporate information about the spread of a component in relation to the covered brain volume when deciding how broad the spatial extent of a component is. Furthermore, anatomical information still needs consideration. A source could appear statistically broad but show a clear peak in cortical gray matter, whereas all other weights are on electrodes in the surrounding white matter; accordingly, one might still consider this source as a local one. ICA can also help to identify which signal an electrode is picking up (compare Fig. 2). An electrode that is located between two neighboring structures could pick up the same source as other electrodes that are located in only one of the structures. This way, the statistical dependencies between channels can be informative of electrode location, when the anatomical information is ambiguous.

A more general criticism about employing ICA could be that components may not always be local and could be hard to interpret. This may be the case when distant sources are correlated because there is communication between the regions. In practice, the filters would still extract time courses that capture the linear dependencies between the distant sources and if the components are of interest, projecting them back could resolve ambiguities in their interpretation. Another potential solution to this would be to apply ICA separately to different electrode shafts. Regardless of the complications one might experience with the exact interpretation of independent components, it is at least possible to inspect how a linear superposition of time courses mixes into the observed data at hand. Together with prior knowledge about the anatomical locations of electrode contacts and the type of surrounding tissue, ICA therefore allows for an informed decision about anatomical sources of recorded activity. Filtering the data with coefficients that were derived via ICA can therefore be a crucial advantage in determining anatomical sources of effects and it is a more informative preprocessing choice then the application of a fixed predetermined re-referencing scheme which does not account for the complex dependencies that may exist between channels.

## Acknowledgements

This work was supported by grants from the Deutsche Forschungsgemeinschaft (HA 5622/1-1), the European Research Council (Grant Agreement no 647954) awarded to SH, and a Sir Henry Dale Fellowship jointly funded by the Wellcome Trust and the Royal Society (107672/Z/15/Z) awarded to BS. SH is further supported by the Wolfson Society and the Royal Society. CK is supported by the Stiftelsen Olle Engkvist Byggmästare.

